# Idiosyncratic liver alterations of five frog species to land use changes in the Brazilian Cerrado

**DOI:** 10.1101/837534

**Authors:** Lilian Franco-Belussi, Diogo B. Provete, Rinneu Elias Borges, Classius de Oliveira, Lia Raquel de Souza Santos

## Abstract

Changes in land use trigger environmental changes that can lead to decreased biodiversity and species loss. The liver is an essential detoxification organ that reflects systemic physiological responses to environmental changes. Here, we tested whether land use changes influence the amount of substances from the hepatic cellular catabolism and melanomacrophages of five anuran species in the Brazilian Cerrado. We used routine histological and histochemical techniques. We then use recently developed ecological methods to relate functional traits to environmental variables. There was an increase in the amount of melanin in environments with high proportion of agriculture, as well as variation in the amount of lipofuscin and hemosiderin. Therefore, the area of melanomacrophages in the liver and the metabolic products in their cytoplasm can be used as biomarkers of environmental changes in regions with intense agricultural activities. Our results add a new perspective to the influence of land use changes on environmental health by highlighting the effect of environmental changes on internal morphological aspects of animals.

## 1. Introduction

Human-driven land use changes are causing biodiversity loss around the world (Powers and Jetz, 2019). Brazil is one of the countries with the highest proportion of net loss of tree cover in South America, with a loss of 8% from 1982 to 2016 (Song et al., 2018). At the same time, there was a 12% increase in short vegetation cover (Song et al., 2018), which includes shrubs and herbaceous vegetation. This trend was especially significant in the Brazilian Cerrado, of which less of 2% are protected (Françoso et al., 2015; Damasco et al., 2018). Accordingly, more than half of the original 2 million km^2^ of the Cerrado were transformed into planted pastures and annual cultures by 2005 (Klink and Machado, 2005). Central Brazil is a thriving region for industrial agricultural activities as one of the largest exporters of soybean and cattle meat in the world (Contini and Martha 2010, Reynolds et al., 2016). One of the consequences of land use change for export-oriented agricultural activities is the intensive use of agrochemicals (Schiesari et al., 2013; Aranha and Rocha, 2019). Therefore, land use changes, along with agrochemicals, are currently the main threats for biodiversity conservation and the sustainable use of natural resources in this biome (Reynolds et al., 2016).

Water quality in the Cerrado has been drastically affected by the intense use of fertilizers, with a significant increase in nitrogen and pesticides (Hunke et al., 2014). As a result, freshwater habitats in the Cerrado receive a great load of contaminants, which impact several aspects of aquatic biodiversity (De Marco et al., 2013; Bichsel et al., 2016). For example, there is evidence that the replacement of natural habitats by urban and agricultural areas decrease not only amphibian populations, but also their genetic diversity (Eterovick et al., 2016). Additionally, land use changes can promote rapid transformation in biological communities beyond species composition. Specifically, it can alter phenotypic aspects of several animal groups (e.g., Borges et al., 2019a), which impact how species interact and adapt to a changing environment. These phenotypic changes include DNA damages (Borges et al., 2019b) and developmental abnormalities (Borges et al., 2019a). However, little is known about the effects of land use changes in internal phenotypic aspects of organisms inhabiting Neotropical agroecosystems.

Amphibians are good bioindicators of environmental quality because they show rapid responses to environmental stress (Brodeur and Candioti, 2017). In addition, they have permeable skin and eggs, making them vulnerable to aquatic contamination and infections. As such, these animals are useful for monitoring changes in both the aquatic and terrestrial environments because they depend on the two environments to complete their life cycle (Brodeur and Candioti, 2017). Amphibians are rapidly declining worldwide due to human-induced changes in the environment (Catenazzi, 2015; Alton and Franklin, 2017). Several factors are involved in this decline, including climate change, increased incidence of ultraviolet (UV) radiation due to ozone depletion, invasive species, environmental contamination, diseases, and habitat change or loss (Collins and Crump, 2009; Alton and Franklin, 2017).

Previous studies have analyzed the effects of agrochemicals on tadpole developmental abnormalities (Borges et al., 2019a) or genotoxic effects in adult anurans. However, environmental alterations that promote morphological and physiological changes at the tissue level are poorly understood. The liver plays a key role in the detoxification of xenobiotics (Pérez-Iglesias, et al., 2018; Fanali et al., 2018), as well as other functions related to the metabolism (Hinton et al., 2001; Fenoglio et al., 2005). The detoxification is performed by hepatocytes and melanomacrophages and may either enzymatic or not. Melanomacrophage centers and their pigments (melanin, lipofuscin, and hemosiderin) are involved in the hepatic response to various toxic compounds. Thus, the liver is an important organ to evaluate the response of organisms to environmental pollutants (Fenoglio et al., 2005).

Melanomacrophages (MMs) are cells that occur in the hepatic tissue and produce and store melanin in their cytoplasm. The area and occurrence of melanomacrophages in the liver increase in response to environmental stressors, such as temperature (Santos et al., 2014), UV radiation (Franco-Belussi et al., 2016), and xenobiotics (Pérez-Iglesias et al., 2016). In addition to melanin, MMs contain catabolic substances originated from the degradation of red blood cells: hemosiderin and lipofuscin, originated from the degradation of polyunsaturated membrane lipids (Oliveira and Franco-Belussi 2012). Therefore, the density of melanomacrophages can be a useful morphological biomarker for the effect of environmental stressors (Oliveira et al., 2017). Morphofunctional changes in the hepatic parenchyma happen as a result of contaminants, suggesting the action of detoxification by melanomacrophages due to the processing action of some enzymes, besides the protective action of melanin (Fenoglio et al., 2005). Additionally, the functions of melanomacrophages are also related to the maintenance of homeostasis, regulating fibrosis, controlling basophiles and participating in the recycling of red blood cells (Gutierre et al. 2017).

Changes in the amount of catabolic substances within melanomacrophages may be associated with changes in phagocytic activity (Bucke et al., 1992; Fenoglio et al., 2005). For example, lipofuscin is produced as a result of the degradation of cellular components. Thus, its increase may hinder cell renewal and promote an accumulation of damaged cellular components in the tissue. The accumulation of lipofuscin can increase cellular sensitiveness to oxidative stress, since this molecule binds to metals, such as iron and copper (Terman and Brunk, 2004). Hemosiderin is an intermediate metabolite generated during the recycling of components in blood production. Thus, the accumulation of hemosiderin indicates a disorder in blood cell recycling (Pérez-Iglesias et al., 2016). In addition, environmental factors that alter the concentration of these substances in the liver may contribute strongly to the decrease in animal health. Consequently, this can affect individual fitness and promote population decline. Therefore, analyzing liver cell physiology may be useful for detecting the effects of environmental changes by human actions. However, while previous studies (Franco-Belussi et al. 2013, Santos et al. 2014, Gregorio et al. 2016, Pérez-Iglesias et al. 2016) have already analyzed the variation of melanin, lipofuscin, and hemosiderin in response to climatic factors and xenobiotics in laboratory experiments using model species, no study has investigated how these three substances vary at the same time in response to changes in land use patterns in wild amphibian populations.

Here, we tested the influence of land use changes associated with aquatic contaminants on liver cell morphology and physiology of five frog species. These can be hidden effects of changes associated with agricultural intensification that are often neglected in biodiversity assessments that only consider species abundance and occurrence. Aquatic contaminants may alter organismal health that cannot be assessed by only recording species presence in each environment, since the internal morphology of individuals can be damaged, which can compromise population viability and species persistence in the environment in the long term.

## 2. Material and methods

### 2.1. Study area and specimen sampling

Field work was carried out in two regions: three ponds in the surroundings of Rio Verde (17°48’11.28” S; 50°56’24.95” W), and two ponds within the Emas National Park (18°15’32.11” S; 52°53’14.13” W; Fig. 1), Goiás, central Brazil. Samplings were conducted during the breeding season between November and March for three consecutive years: 2013, 2014, and 2015. The region around the city of Rio Verde is a thriving agricultural area, mainly covered with planted pastures, and monocultures of soybean, corn, and sorghum. The Emas National Park is an enclave of Cerrado Protected Area with several typical vegetation types of this biome, varying from open formation, such as *campo limpo* and *campo rupestre*, to closed-canopy formations, such as *Cerradão* and Seasonal Deciduous Forest (Valente 2006).

**Fig. 1.**
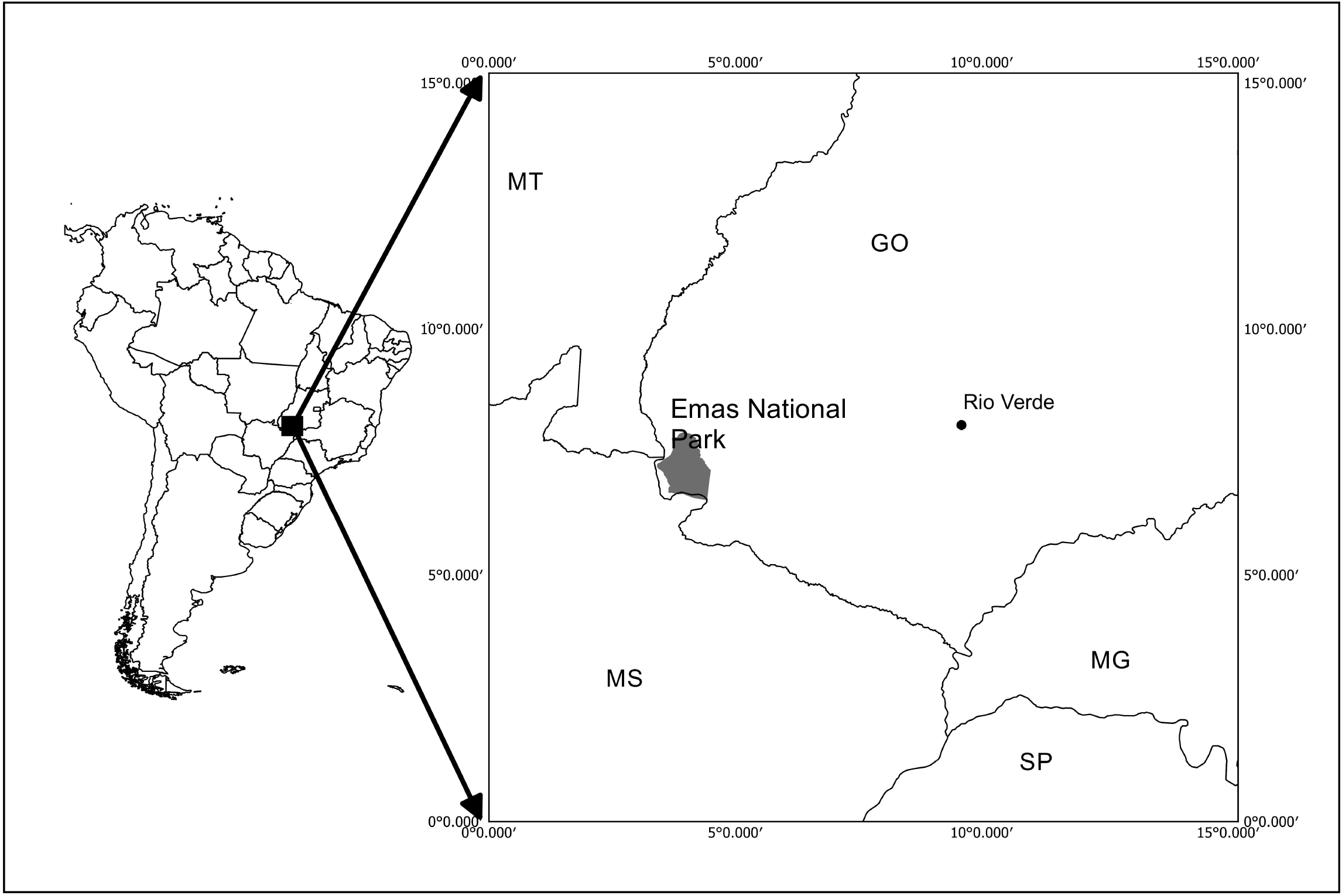
Map showing the location of sampling sites in Central Brazil.

We used a paired design in which we collected five adult males of five frog species in both regions: *Boana albopunctata, Dendropsophus minutus, Scinax fuscomarginatus, Leptodactylus fuscus*, and *Physalaemus cuvieri*. These species were used because they occurred in both regions. Additionally, all of them are widely distributed throughout South America (AmphibiaWeb, 2019) and seem to be generalists, occurring in a wide range of habitats, from open, natural formations, to bush Savannas, to peri-urban areas (Haddad et al. 2015). *Boana albopunctata, Scinax fuscomarginatus*, and *Dendropsophus minutus* are hylid tree-frogs that call perched on the vegetation, while *Leptodactylus fuscus* and *Physalaemus cuvieri* are medium-sized leptodactylids that are found calling and usually foraging on the ground near temporary ponds (Haddad et al. 2015). Both *B. albopunctata* and *D. minutus* deposit egg masses directly on water, while *L. fuscus* and *P. cuvieri* build foam nests in which they deposit eggs on the margin (in the case of *P. cuvieri*) or on subterranean chambers (in the case of *L. fuscus*; Haddad et al. 2015) of water bodies. Thus, adults, eggs, and larvae of these species have different degrees of contact with contaminated water. Consequently, these species can potentially be useful as environmental indicators, since they are highly abundant within their geographic range and seem to respond quickly to environmental disturbance.

### 2.2. Water quality analysis

To test if the ponds used by amphibians were contaminated by pesticides, we collected 1-L samples of water in one pond from each region. We quantified organochlorines and organophosphates in water samples from both sites. Quantification was made using standard methodology by A3Q Laboratory of Environmental Quality (Cascavel, Paraná, Brazil). Samplings from the Park did not contain any substance above the references, while Rio Verde had atrazine well above the level allowed by the Brazilian environmental agency (Table S1).

### 2.3. Routine histological processing

Specimens were anesthetized with 5 g/L benzocaine. This procedure was approved by the ethics committee on animal use of our university (protocol #0316, CEUA/UniRV). Liver fragments were extracted and fixed in metacarn solution for 3 h. Subsequently, they were dehydrated in an alcoholic series and included in historesin (Leica^®^). We obtained 2 μm sections in rotating microtome (RM 2265, Leica, Switzerland), which were stained with hematoxylin and eosin.

Histochemical analyzes were performed for the detection of lipofuscin as follows: sections were incubated for 15 min in Schmorl’s solution, composed of 75 mL of 1% ferric chloride, 10 mL of potassium ferricyanide, and 15 mL of distilled water. Then, sections were immersed in neutral red aqueous solution at 1% followed by 1% aqueous solution of eosin. For the detection of hemosiderin, sections were incubated for 15 min in acid solution of ferrocyanide obtained by the dissolution of 2 g of potassium ferrocyanide in 100 mL of 0.75 mol/L hydrochloric acid solution. Subsequently they were immersed in aqueous solution of 1% neutral red followed by aqueous solution of 1% eosin.

Images were captured in a digital camera coupled with a microscope (Laborana Lab50AB-S) using an image capture system (Lab3001-C) and analyzed in Image Pro-Plus v. 6.0 (Media-Cybernetics Inc.). Image analysis was conducted using 25 randomized histological fields for each animal. Specifically, we quantified the area occupied by each substance by differences in staining intensity in 25 pictures per animal following Santos et al. (2014).

### 2.4. Quantification of land use pattern

To quantify land use, we used a shape file with land cover and use classes (IBGE, 2018 URL: http://www.sieg.go.gov.br/produtosIMB.asp?cod=4725). This file classifies land use and cover into 14 classes based on satellite images from 2014 (see Appendix II in IBGE 2018). To calculate the area of each land use, we established a buffer of 500 m radius around each water body sampled and calculated the area occupied by each land use class in ArcGIS 9.0 software (ESRI 2011). This buffer size has been commonly used in studies of landscape ecology involving anurans (e.g., Almeida-Gomes et al. 2016), since it is usually the dispersal distance of individuals moving among ponds in agricultural landscapes. The areas of each land use class were then converted into proportions and used as predictor variables in further analyzes. The land use types we found around our sampling sites were: natural grassland and shrub vegetation, artificial area, forest mosaic, grassland mosaic, planted pastures, and natural pasture, and agricultural area. Since the area of some land use types in our data set was small, we combined artificial area with grassland mosaic into a class called anthropized area, and also combined forest mosaic, natural pastures, and agricultural area into a class called farming to improve data analysis and interpretation of results.

### 2.5. Data analysis

There are currently several methods to test the influence of environmental variables on species traits (see review in Kleyer et al., 2012), the so-called fourth-corner problem. However, there is still no consensus as to which method is best, and it seems that this decision is dependent on the context and trait data type (continuous, categorical or mixed), although studies show that correlating Community-weighted Mean with environmental variables always seems to produce larger Type I Errors (Peres-Neto et al., 2017; ter Braak et al., 2017). Here, we used a double-constrained Correspondence Analysis (or dc-CA, for short) to test whether the mean area of melanin, hemosiderin, and lipofuscin (response variables) of the five species are correlated with different classes of land use (predictor variables). This is a long-proposed method (Lebreton et al., 1988), but a new algorithm has only recently been proposed (ter Braak et al., 2018). The advantages of dc-CA are that it considers the correlation among environmental variables and among traits, when relating traits to environment (see ter Braak et al., 2018), differently from RLQ and CWM-RDA (see Kleyer et al. 2012). dc-CA uses three tables: a species-by-site matrix **Y**, which can contain either abundances or incidence; a site-by-environment matrix **E**, and a species-by-trait **T** matrix. The method starts by conducting a Correspondence Analysis constraining both the columns (species) by species traits and row scores of the **Y** matrix by environmental variables (ter Braak et al. 2018). We then produced a triplot showing the relationship among traits, environmental variables, and species incidence in a single ordination diagram. Analysis was conducted using R code available in the supplementary material of ter Braak et al. (2018) and Canoco 5.12 (ter Braak and S□milauer 2018).

## 3. RESULTS

We found both great intra- and interspecific variation in the amount of each substance (Fig. 2). Most species had higher amounts of lipofuscin and melanin than hemosiderin, except *Boana albopunctata* that had lower amounts of lipofuscin than the other species. Interestingly, *Leptodactylus fuscus* and *Dendropsophus minutus* showed little intraspecific variation in the amount of all three substances, while *Physalaemus cuvieri* and *B. albopunctata* had high intraspecific variation. The means for the three substances was similar for *L. fuscus* and *D. minutus*, whereas the other three species had very distinct means.

**Fig. 2.**
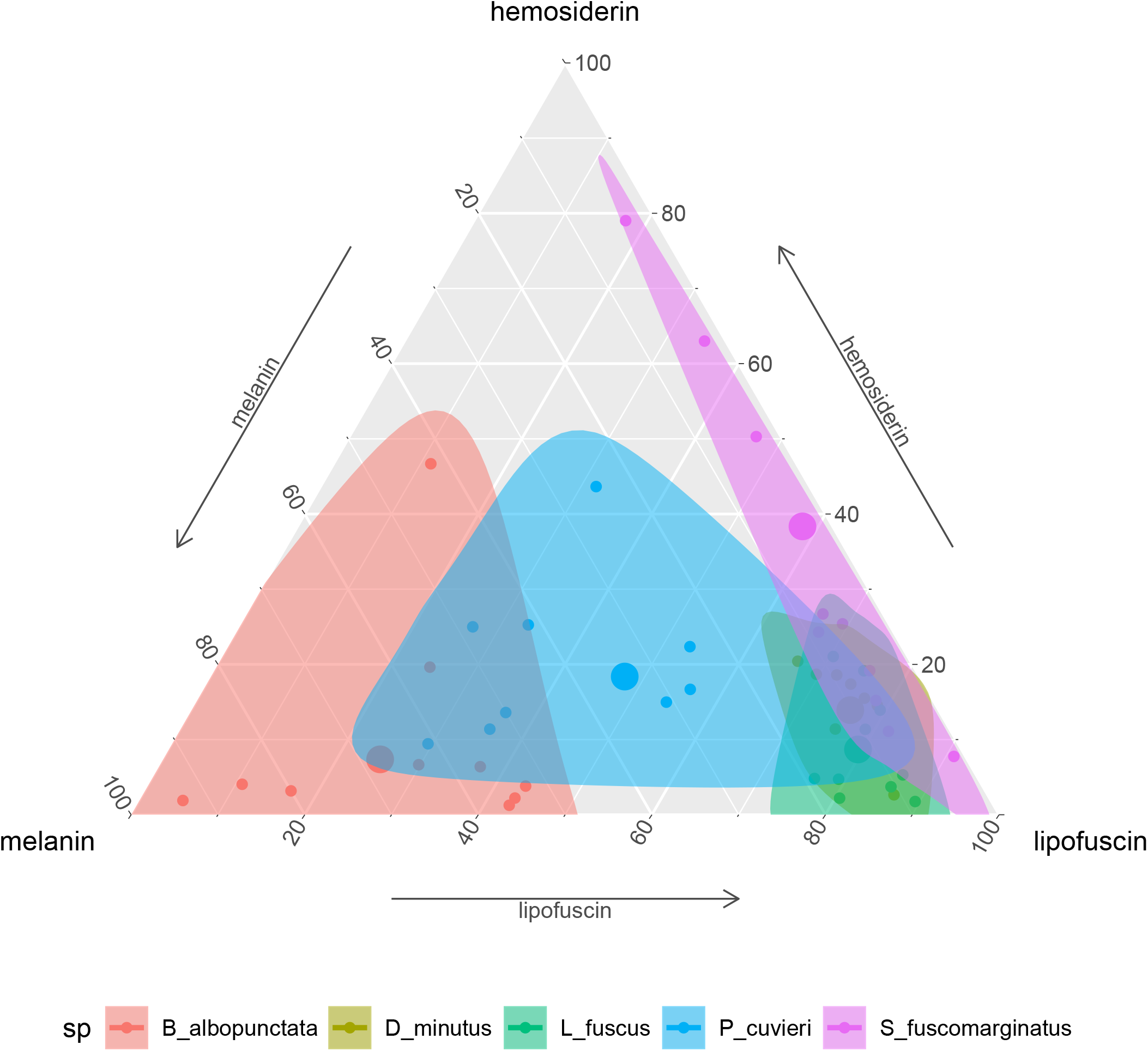
Ternary plot showing the relative proportion of the area of each substance in the five frog species. Small points indicate individual measurements, while the large dot represents the mean of each substance for each species.

The total inertia of the dc-CA model was 0.955. The first axis of dc-CA accounted for 72% of the variation in the trait-environment relationship, while the second accounted for 26%. The maximized fourth-corner correlation along the first and second axes are 0.83 and 0.50, respectively. Melanin was positively correlated with natural and planted pastures, but negatively correlated with man-made buildings and agriculture (Table 1). Hemosiderin was negatively correlated natural and planted pastures, and man-made buildings, but positively correlated with agriculture. The correlation pattern of lipofuscin was almost identical to hemosiderin with small changes in the strength of correlation with some land use types (Table 1).

**Table 1.** Fourth-corner correlation calculated with between species traits (area of each substance in liver) and environmental variables (percentage of land use type in 500 m buffer).

|  | Melanin | Hemosiderin | Lipofuscin |
| --- | --- | --- | --- |
| Natural pasture | 0.139 | -0.276 | -0.252 |
| Man-made building area | -0.033 | -0.188 | -0.228 |
| Planted pastures | 0.473 | -0.135 | -0.267 |
| Agriculture | -0.386 | 0.721 | 0.78 |

The occurrence of *P. cuvieri* was positively correlated with the percentage of planted pasture in the landscape (Fig. 3). This was also the species with the highest amount of melanin in the liver (Fig 3), which was positively correlated (fourth-corner correlation = 0.473, Table 1) with the percentage of planted pasture (Fig. 3). In contrast, *L. fuscus* had the lowest amount of melanin (Fig. 3) whose incidence was negatively correlated with the percentage of planted pasture in the landscape (Fig. 3).

**Fig. 3.**
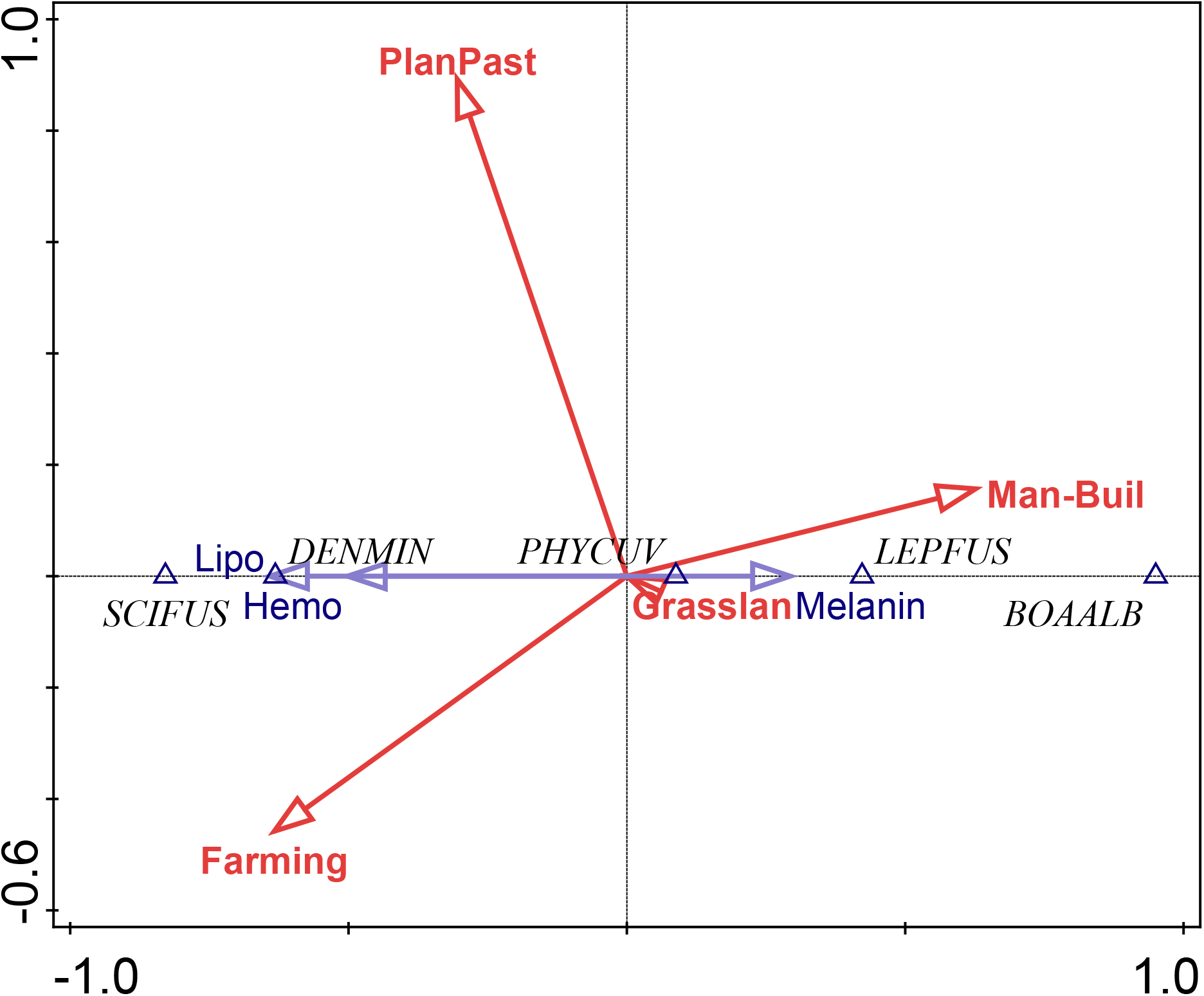
Ordination diagram of double-constrained Correspondence Analysis showing the relationships between species incidence, cell metabolism substances, and land use classes. Arrow length indicates the importance of variables for the construction of ordination axes.

Conversely, the incidence of *D. minutus* and *S. fuscomarginatus* were positively correlated with area of agriculture and negatively with anthropogenic area, while the incidence of *B. albopunctata* was positively correlated with anthropogenic area and negatively with area of agriculture (Fig. 3). *Dendropsophus minutus* and *S. fuscomarginatus* had higher amounts of hemosiderin and lipofuscin, suggesting higher hepatic metabolism, which was positively correlated with agriculture and negatively with percentage of anthropogenic area (Fig 3). Conversely, *B. albopunctata* had fewer hepatic catabolite substances (Fig. 3).

## 4. Discussion

We found that *P. cuvieri* occurred in sites with planted pastures and man-made buildings, had higher amounts of melanin, while the occurrence of *L. fuscus* was positively correlated with man-made buildings and negatively correlated with planted pastures. Additionally, *D. minutus* and *S. fuscomarginatus* had high amount, while *B. albopunctata* had low amount of lipofuscin and hemosiderin and was positively correlated with man-made buildings. Taken together, these results demonstrate the effects environmental changes on melanomacrophages and hepatic metabolism of these frog species.

The amount of melanin was highest in *P. cuvieri* and *B. albopunctata*, but low in *L. fuscus, D. minutus*, and *S. fuscomarginatus*. There was also a high positive correlation between melanin and planted pastures (0.473), and a less strong correlation with man-made buildings (−0.033), and natural pastures (0.139). Melanin is a complex polymer that absorbs and neutralizes free radicals, besides participating in the innate immune response and protection of tissues in ectotherms (Cesarini 1996). Changes in the amount of melanin was found in frogs experimentally exposed to several xenobiotics. In these experiments, the variation in melanin was due to several compounds (reviewed in Oliveira et al. 2017), such as polycyclic aromatic hydrocarbons (PAHs; Fanali et al. 2018), herbicides (e.g., atrazine and glyphosate; Pérez-Iglesias et al. 2019; Bach et al. 2019), drugs, and endocrine disrupters (Gregorio et al. 2016). Here, we found large amounts of atrazine in the area under agriculture influence (more than 5,000 time the limit accepted by Brazilian legislation, i.e. 5349.940 μg/L). Atrazine is an endocrine disruptor, which has immunotoxic and immunosuppressive effects even at concentrations usually found in the environment (i.e. < 500 μg/L; Rohr ad McCoy 2010). This substance can change swimming ability, cause malformations, and promote nuclear alterations at higher concentrations in tadpoles (Pérez-Iglesias et al. 2019). Atrazine causes oxidative stress in several tissues, which can evolve to function loss.

Although our statistical model explained much variation in the three substances, other climatic factors that have been changing as a consequence of human activities, such as temperature and Ultra-violet radiation can also change the amount of melanin in internal tissues of frogs (Franco-Belussi et al. 2017). Changes in these environmental variables may promote changes in hepatic metabolism, such as increasing glycogen and lipid levels (Mizell, 1965; Barni & Bernocchi, 1991; Fenoglio et al. 1992, Barni et al. 2002). Adaptation to natural conditions (i.e., hibernation) may promote an increase in the amount of melanin in the liver by increasing the number of cells (i.e. hyperplasia) or the increase in cell size (i.e. hypertrophy), besides an increase in the production of melanin in cell interior (Barni et al. 2002). These changes occur as a mechanism of metabolic defense against free radicals. For example, with increased storage of lipids and glycogen in hepatic tissue due to environmental changes that occur naturally in hibernating animals (Mizell, 1965; Barni & Bernocchi, 1991; Fenoglio et al. 1992, Barni et al. 2002). Thus, the liver of anurans is a plastic organ, besides being sensitive to the alterations of the natural biological rhythms (i.e. reproduction and hibernation), coordinating these mechanisms to maintain the homeostasis of the organism during adaptation to environmental changes (Barni et al. 2002). Therefore, cells that produce and store melanin seem to be involved in the adaptation to environmental stressors at the tissue level.

*Dendropsophus minutus* and *S. fuscomarginatus* had high amounts, whereas *B. albopunctata* had low amounts of hemosiderin and lipofuscin. Lipofuscin and hemosiderin are products of cellular catabolism and can be used to measure hepatic metabolism, since both substances may be altered as a result of environmental stress following habitat alteration (Santos et al. 2014, Pérez-Iglesias et al. 2016). Low amounts of these substances indicate decreased phagocytic activity of cells (Bucke et al. 1992; Fenoglio et al. 2005), while accumulation of hemosiderin is related to increased turnover of blood cells (Fenoglio et al. 2005). Therefore, the increase in recycling of blood cells in *D. minutus* and *S. fuscomarginatus* indicates that biotic or abiotic factors resulting from anthropic changes may be causing changes in the liver of these two species, but so in *B. albopunctata*.

The idiosyncratic responses of species may be related to differences in life history traits. For example, *Physalaemus* and *Leptodactylus* are terrestrial and can have more contact with xenobiotics; while *Boana, Scinax*, and *Dendropsophus* are arboreal and putatively less exposed to aquatic xenobiotics (Silva et al. 2013). However, it is noteworthy that any drastic change in land use appears to promote metabolic alterations related to liver physiological processes in anurans. Therefore, MMCs seem to be an efficient biomarker that indicated alterations in the liver in response to land use change. Actions for mitigating the negative effects of industrial agricultural activities must be taken if we want to reach the goal of making environmentally responsible agricultural products (ONU 2050 goals), especially in savanna biomes (Parr et al. 2014, Overback et al. 2015).

## Acknowledgements

Funding: This study was supported by Conselho Nacional de Desenvolvimento Científico e Tecnológico (CNPq) (grant #477044/2013-1) to L.R.S.S and Fundação de Amparo a Pesquisa do Estado de São Paulo (FAPESP) (grant #2013/02067-5). LFB was supported by a FAPESP post-doctoral fellowship (#2014/00946-4) during the initial phase of this study. B. Valverde, W. R. Rezende, R. A. Assis, and G. Leite helped with histological procedures. R. F. Oliveira in the elaboration of the map with the animal collection points. M. Almeida-Gomes helped with GIS data processing and extracting area of each land use type. C. ter Braak clarified some statistical aspects of analysis. CO has been continuously supported by CNPq (#305081/2015-2). The Instituto Chico Mendes de Conservação da Biodiversidade provided colleting permit (Sisbio #34485-1) and Emas National Park provided housing and authorization for field work.

