## Supplementary material for "Idiosyncratic liver alterations of five frog species to land use changes in the Brazilian Cerrado": Table S1

| **Agrochemicals** | **Concentration (µg/L)** | | **Reference value (µg/L)**^1^ |
| --- | --- | --- | --- |
|  | Emas National Park | Rio Verde |  |
| 2.4-D + 2.4.5-T | <0.01 | <10.0 | 30.0 |
| Aldicabe+Aldicarbe-sulfone+Aldicarbe-sulfoxide | <0.01 | <5.0 | 10.0 |
| Alachlor | <0.05 | <0.053 | 20.0 |
| Aldrin | <0.03 | <0.03 | 0.03 |
| Dieldrin | <0.03 | <0.03 | 0.03 |
| Atrazine | <0.05 | 5349.94 | 2.00 |
| Carbendazim+Benomyl | <0.01 | <20.0 | 120.0 |
| Carbofuran | <0.10 | <1.048 | 7.00 |
| Chlordane (cis+trans) | <0.01 | <0.011 | 0.20 |
| Chlorpyrifos+ Chlorpyrifos-Oxon | <0.10 | <0.105 | 30.0 |
| DDT (4.4-DDT+4.4-DDE+4.4DDT) | <0.01 | <0.011 | 1.00 |
| Diuron | <0.01 | <20.0 | 90.0 |
| Endossulfan (alpha+beta+sulfate) | <0.10 | <0.101 | 20.0 |
| Endrin | <0.03 | <0.031 | 0.60 |
| Glyphosate+AMPA | <0.01 | <110.0 | 500.0 |
| Lindane | <0.01 | <0.01 | 2.00 |
| Mancozeb | <0.01 | <20.0 | 180.0 |
| Metolachlor | <0.05 | <0.050 | 10.0 |
| Methamidophos | <0.01 | <1.00 | 12.0 |
| Molinate | <0.10 | <0.104 | 6.00 |
| Metil Parathion | <0.25 | <0.250 | 9.00 |
| Pendimenthalin | <0.05 | <0.051 | 20.0 |
| Permethrin | <0.10 | <0.103 | 20.0 |
| Profenofos | <0.01 | <20.0 | 60.0 |
| Simazinee | <0.05 | <0.051 | 2.00 |
| Tebuconazole | <0.01 | <20.0 | 180.0 |
| Terbufos | <0.01 | <0.5 | 1.20 |
| Trifluralin | <0.05 | <0.051 | 20.0 |

^1^ In accordance to Brazilian Enviromental Comittee (Portaria nº 2.914 de 12 de Dezembro de 2011).
